# CryoEM Reconstruction of Yeast ADP-Actin Filament at 2.5 Å resolution. A comparison with mammalian and avian F-actin

**DOI:** 10.1101/2024.09.13.612689

**Authors:** Sarah R. Stevenson, Svetomir B. Tzokov, Indrajit Lahiri, Kathryn R. Ayscough, Per A. Bullough

## Abstract

The core component of the actin cytoskeleton is the globular protein G-actin, which reversibly polymerises into filaments (F-actin). Budding yeast possesses a single actin which shares 87-89% sequence identity with vertebrate actin isoforms. Previous structural studies indicate very close overlap of main-chain backbones. Intriguingly however, substitution of yeast *ACT1* with vertebrate β-cytoplasmic actin severely disrupts cell function and substitution with a skeletal muscle isoform is lethal. Here we report a 2.5 Å structure of budding yeast F-actin. Previously unresolved side-chain information now highlights four main differences in the comparison of yeast and vertebrate ADP F-actins: a more open nucleotide binding pocket; a more solvent exposed C-terminus; a rearrangement of intersubunit binding interactions in the vicinity of the D-loop and changes in the hydrogen bonding network in the vicinity of histidine 73 (yeast actin) and methyl-histidine 73 (vertebrate actin).

## INTRODUCTION

The actin cytoskeleton is a ubiquitous feature of eukaryotic cells capable of performing diverse cellular functions in response to the needs and specialisms of the cell. The main component of the actin cytoskeleton is the globular protein actin (G-actin), which reversibly polymerises into filaments (F-actin). These monomeric and filamentous forms exist in cells in a dynamic and highly regulated equilibrium.

The actin monomer is a highly conserved ∼42 kDa globular protein of around 375 amino acids. The protein fold has been categorized into four subdomains (SD1-SD4; Figure 1A,B), with the N and C-termini both located in subdomain 1. Critical for function is its ATPase activity, with each actin monomer bound to one nucleotide in a deep ATP-binding cleft that separates the protein into two major globular domains^1^. Binding of ATP, or its hydrolysed form ADP, is stabilized by a divalent cation in the cleft near the terminal phosphate of the nucleotide. The hydrolysis of ATP occurs rapidly upon polymerisation, but release of the γ-phosphate (P_i_) is not immediate, resulting in ADP-P_i_ in the nucleotide cleft along a segment of the filament. ATP hydrolysis is irreversible, and nucleotide exchange only occurs in the monomeric actin form^2^.

**Figure 1.**
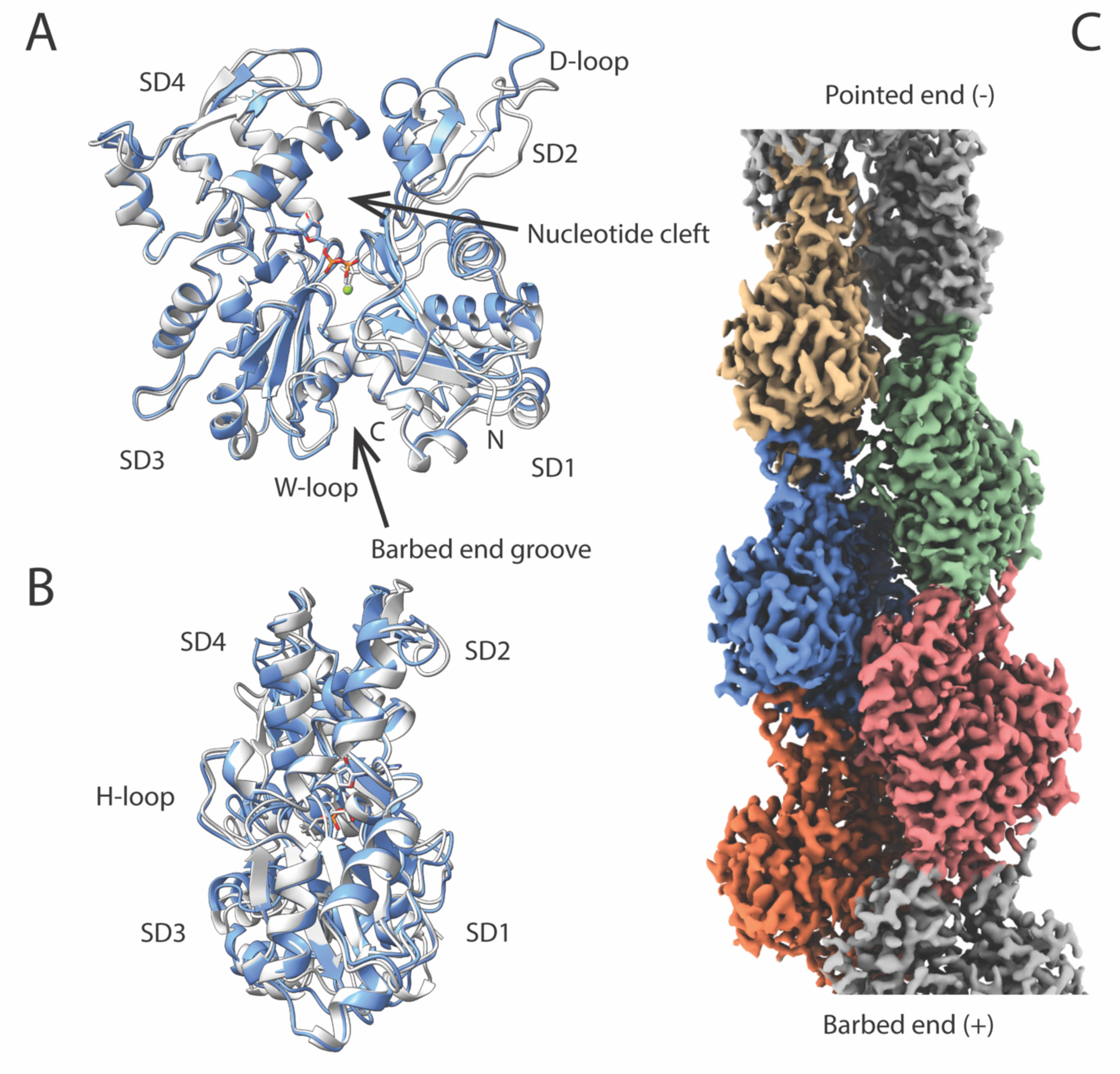
CryoEM structure of yeast F-actin at 2.5 Å resolution. (A) Backbone fold of a single yeast F-actin protomer (blue) compared with yeast G-actin derived from the structure of a yeast G actin-gelsolin complex^7^ (gray) (PDB ID:1YAG). Subdomains 1-4 are labeled SD1-SD4. The ADP is shown in stick form along with the Mg^2+^ (green). (B) View of the protomer rotated approximately 90° from (A). (C) CryoEM reconstruction of a segment of the F-actin filament. 5 protomers are shown in color. (See also Figs. S1,S2).

Within an actin filament, the helically arranged actin monomers are all oriented in the same direction, conferring an overall polarity (Fig. 1C). As a result of this polarity, the two ends of an actin filament have different properties, most notably different rates of polymerisation, with a fast-growing ‘barbed’ end and a slow growing ‘pointed’ end (also referred to as plus and minus ends respectively). Because one end of the filament elongates more rapidly than the other, this results in a gradient of bound nucleotide along the length of the filament, with ADP in the cleft of the oldest regions of the filament; ADP-P_i_ in newer sections; and a small number of ATP-bound monomers at the actively growing filament end^3,4^.

At a structural level, on incorporation into a filament, an actin monomer undergoes a conformational change that can be thought of as flattening. This is achieved by the relative rotation of the two major domains of the protein found on either side of the cleft (SD1+SD2, and SD3+SD4) by approximately 20° (Fig. 1A,B)^5^. This conformational change activates ATP hydrolysis by a subtle reorganization of the active site that increases its catalytic activity by several-thousand fold^6^. A key factor in this increased catalytic activity is the repositioning of Gln137 and His161 relative to the γ-phosphate of ATP^5^. This is because these residues each anchor a water molecule that plays an active role in the nucleophilic attack on the γ-phosphate during ATP hydrolysis^7,8^.

On the opposite side of the monomer to the nucleotide binding cleft is a shallow barbed end groove between SD1 and SD3 that is the binding site for several actin binding proteins^9,10^. The groove includes a region referred to as the W-loop (residues 165-172). Other distinct structural features of the protein backbone are the H-loop (residues ∼262-272, located between subdomains 3 and 4) and the DNase I binding loop (D-loop; residues 40-50 in subdomain 2, (Figure 1A,B). The D-loop, H-loop and N-terminus of G-actin are highly mobile, and remain so in the filament^11^. As well as flattening of the molecule, another notable change upon polymerisation is the reconfiguration of the D-loop^5^, which is a major player in inter-filament contacts (Fig. 1A).

The budding yeast *S. cerevisiae* possesses only a single actin isoform, which shares 87-89% sequence identity (94-96% sequence similarity) with vertebrate actin isoforms. X-ray crystallographic studies have also found the monomeric G-actin structures of yeast and vertebrate actin to be highly similar^7^. However, substitution of the yeast gene for the vertebrate β-cytoplasmic isoform severely disrupts cell function^12^, and substitution with the vertebrate skeletal muscle isoform is lethal^13^. Direct comparison of dynamics *in vitro* have indicated that yeast actin nucleates more efficiently than skeletal muscle actin in certain ionic environments, and that yeast F-actin exhibits increased fragmentation compared to skeletal muscle F-actin^14,15,16^. It has also been noted that yeast F-actin does not exhibit the Mg^2+^-dependent stiffness/rigidity of skeletal muscle actin^17^. This has been attributed to a single residue substitution (Glu167Ala) possibly weakening the filament’s interaction with a stiffness-associated cation^18^. Differences have also been observed between yeast and muscle F-actins in their interactions with some binding partners. For instance, yeast F-actin binds phalloidin more rapidly but more weakly than skeletal muscle actin^19^ and has a lower affinity for muscle myosin, which it also activates more weakly^20,21^.

The C-terminal region (subdomain 1) and D-loop (subdomain 2) have also been implicated in functional differences between isoforms. Both these regions exhibit considerable flexibility and are important for intra-strand protomer-protomer interactions. Fluorescent and phosphorescent labeling of Cys374 have indicated that the C-terminus of yeast F-actin is more flexible and more exposed to the surrounding environment than the C-terminus of skeletal muscle F-actin is^22^. The D-loop is susceptible to proteolytic cleavage by the protease subtilisin between Met47 and Gly48 of the D-loop^23^. The rate of subtilisin digestion of yeast F-actin is approximately 10-fold faster than muscle actin, suggested to indicate greater flexibility compared with the D-loop of the skeletal muscle isoform^15^. This has implications for the stability of the intra-strand interactions of yeast F-actin since the D-loop is the major contact site between neighboring protomers within the same strand.

While there are now multiple published structures for skeletal muscle F-actin with resolution of 5 Å or better^4,6,8,11,24,25^ the resolution of yeast F-actin structure has not been reported past ∼20 Å^26,27,28^ until very recently^17^. The key noted differences between earlier density maps and a rabbit skeletal muscle F-actin map of comparable resolution were that yeast F-actin appeared to have reduced inter-strand connectivity as well as a more open nucleotide binding cleft^26^. These observed differences have since been cited as explanations for the biochemical traits of yeast F-actin^22,29^. However, these observations were based on maps in which the only features that could be resolved were individual protomers and the position of the ATP-binding cleft, and therefore require re-investigation with the more advanced technology now available. A more recent paper based on comparisons of ∼4.5 Å resolution structures of wild-type yeast actin and rabbit actin describes apparent differences in the conformation of the D-loop, but these proposed differences also require much higher resolution data to be testable^17^.

The structural information for yeast F-actin obtained by CryoEM shown here is considerably improved in resolution, with detailed information including the position and register of α-helices, side chain conformers and water molecules. Using our 2.5 Å map and an atomic model constructed from it, we investigated whether the formerly reported differences between vertebrate skeletal muscle F-actin and yeast F-actin were consistent with these higher resolution structural data.

## RESULTS AND DISCUSSION

*Saccharomyces cerevisiae* actin (referred to henceforth as yeast actin) was expressed from its single actin-encoding gene *ACT1* in a *Pichia pastoris* expression system according to the method of Hatano and colleagues^30^. In this system actin is expressed and purified as a fusion with the actin monomer binding protein thymosin β4. When this fusion protein is expressed, the thymosin-β4 sequesters the recombinant G-actin, preventing interactions with both the barbed and pointed end of the monomer and also preventing co-polymerisation with *Pichia* actin. Following purification, the actin was cleaved from thymosin-β4 at its normal C-terminal residue using chymotrypsin. Yeast actin was then polymerised for CryoEM analysis as described in Methods.

We reconstructed the yeast actin filaments with Mg^2+^ and ADP (Fig. 1C). We selected 2,047,208 particles from 2,540 micrographs and after several rounds of 2D classification 1,368,572 particles were retained for further processing. These particles were subjected to multiple rounds of 3D helical refinement within CryoSPARC^31^ resulting in a map with a global resolution of 2.5 Å (Fig. S1C-E, Table S1) estimated by Fourier shell correlation with 0.143 criterion (FSC0.143).

The reconstruction clearly showed the ADP and Mg^2+^ well resolved, along with most amino acid side chains and a number of waters (Fig. 2A,D). Densities were weakest (with correspondingly high model B-factors (Fig. S1A)) in the D-loop, peripheral regions of subdomain 4, in the C-terminal region and at the N-terminus, where residues 1 to 5 were not built; nevertheless the full D-loop was still well-resolved in comparison to a number of reported vertebrate F-actin structures (Fig. S1B).

**Figure 2.**
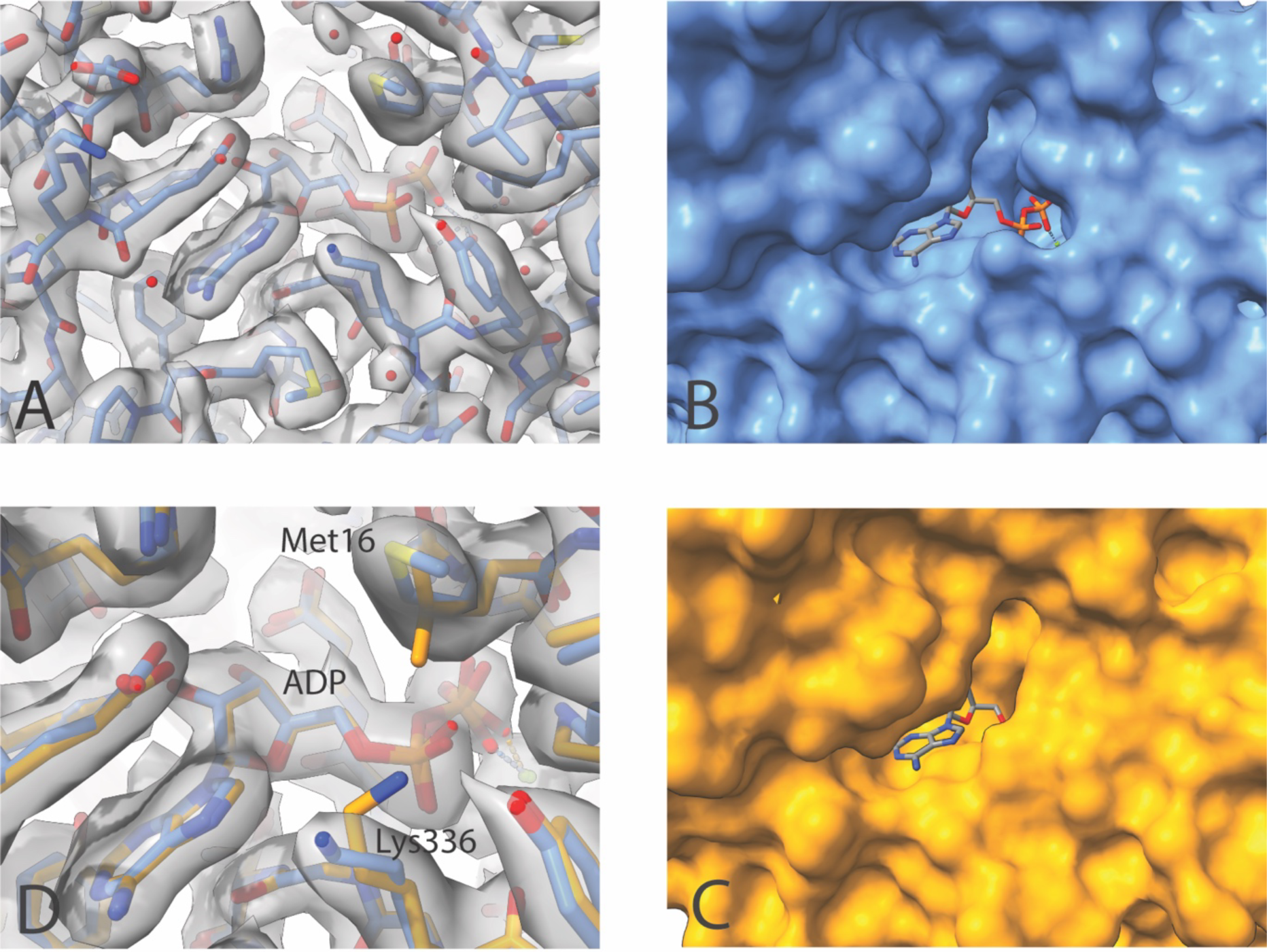
Comparison of the nucleotide binding pocket in yeast and mammalian F-actin. (A) CryoEM density in the vicinity of the binding pocket of yeast F-actin with the fitted atomic model (carbon-cornflower blue; oxygen or water-red; nitrogen-dark blue; sulfur-yellow; phosphorous-orange). (B) Surface rendering of the nucleotide binding pocket in yeast, with the ADP shown in stick form. (C) The equivalent binding pocket in mammalian actin^4^ (PDB ID:8A2T). (D) Comparison of amino acids in the vicinity of the binding site. Yeast carbon-cornflower blue; mammalian carbon-orange. Amino acid labels refer to the yeast actin. (See also Fig. S2).

### Comparison of yeast G and F actin conformations with those of vertebrate actin

With our high resolution structure of yeast F-actin we have the opportunity to make a direct comparison of monomeric and polymerized actin within one species. Comparison of the overall fold of one yeast F-actin subunit and that of yeast G-actin (in complex with gelsolin segment-1^7^) is shown in Fig. 1A. Around the nucleotide binding site we see rearrangements of Gln137 and His161 similar to those reported for avian F-actin with an ATP mimic. However, in our ADP-Mg^2+^ F-actin structure we also see a rotation of the side chain of D154 relative to that in ATP G-actin (Fig. S2A)^6^.

### Comparison with mammalian and avian F-actin CryoEM models

#### Overall architecture

The overall subunit fold is near-identical to that of other recently determined structures from rabbit and chicken (Fig. S3) with backbone rmsd of ∼0.3 to ∼0.6 Å. Within the filament, the axial rise and helical twist between subunits was also similar to rabbit and chicken F-actin at 27.6 Å and -167.2°. We see no significant difference in inter-strand connectivity between yeast and other F-actins. The largest differences in backbone conformation are found in the C-terminal region and the D-loop; it is particularly notable that the C-terminal region in the rabbit ADP-Mg^2+^ bound F-actin determined by Oosterheert *et al.^4^* adopts a significantly different arrangement compared with both yeast actin and other comparable vertebrate actin structures (Fig. S3C). A recently reported comparison of the D-loop conformation of yeast versus rabbit F-actin was based on relatively low resolution (∼4.5 Å) models with very high B-factors (up to 210 Å^2^) in this region; our model (∼2.5 Å resolution) with much lower B-factors (up to 60 Å^2^) reveals the backbone and side-chain rotamers in their more likely position (Fig. S3E)

#### Nucleotide binding site

The nucleotide binding site contains density corresponding to ADP (Fig. 2A) and lacks any density corresponding to a γ-phosphate, indicating this structure represents the ADP state of F-actin. Density for a putative Mg^2+^ ion is also indicated, along with slightly noisier density consistent with the approximate positions of coordinating waters (Fig. S2B). We have not put constraints on the positions of these waters in the model.

Comparison with high resolution structures of avian and mammalian F-actin indicates a more open nucleotide binding cleft in yeast actin (Fig. 2B,C); this is consistent with the observations of Orlova *et al.^26^* on negatively stained fibers. The residues ‘shielding’ the entrance to the skeletal actin site are Leu16 and Lys336 which are in van der Waals contact; in yeast this contact is broken with a Leu16Met substitution and the adoption of a different side chain conformation of Lys336 (Fig. 2D).

His73 is suggested to control the release of P_i_ from actin subsequent to hydrolysis of ATP during polymerization. His73 is a 3-methylhistidine in vertebrate actin^32^ and also in *Plasmodium* actin^33^ but is unmethylated in *S. cerevisiae^34,35^*. The density in our reconstruction is consistent with this. Fig. 3 shows a comparison of residues in the vicinity of His73 for yeast actin and rabbit actin (PDB ID:8A2T). In skeletal F-actin the methylated His73 does not appear to form strong interactions with any neighboring residues whereas in yeast actin the unmethylated His73 is in a slightly more favorable position to form a hydrogen bond with the mainchain oxygen of Gly158. Glu72 is displaced to form a hydrogen bond with Thr77 whilst the slight shift in position of His73, Gly158 and Val159 is accompanied by a shift of the mainchain around Arg177 to Asp179. This prevents a steric clash and leads to a sparser hydrogen bonding network. Arg177 maintains a salt bridge interaction with Asn111 and thus represents the closed ‘gate’ conformation for γ-phosphate release^36^. In our model Oδ2 of Asp179 reaches within 3.8 Å of Nδ1 of His73 (compared with 4.5 Å for 8A2T); given the uncertainty in the map it is possible that Asp179 can adopt a geometry and distance to form a salt bridge with His73, whilst Asp184 appears unlikely to form any interaction^37^. This arrangement is consistent with the “open cleft” conformation examined by Yao *et al.*, although we see no significant difference in the His73 imidazole ring, or salt bridge with Asp184 despite this being previously predicted^38^.

**Figure 3.**
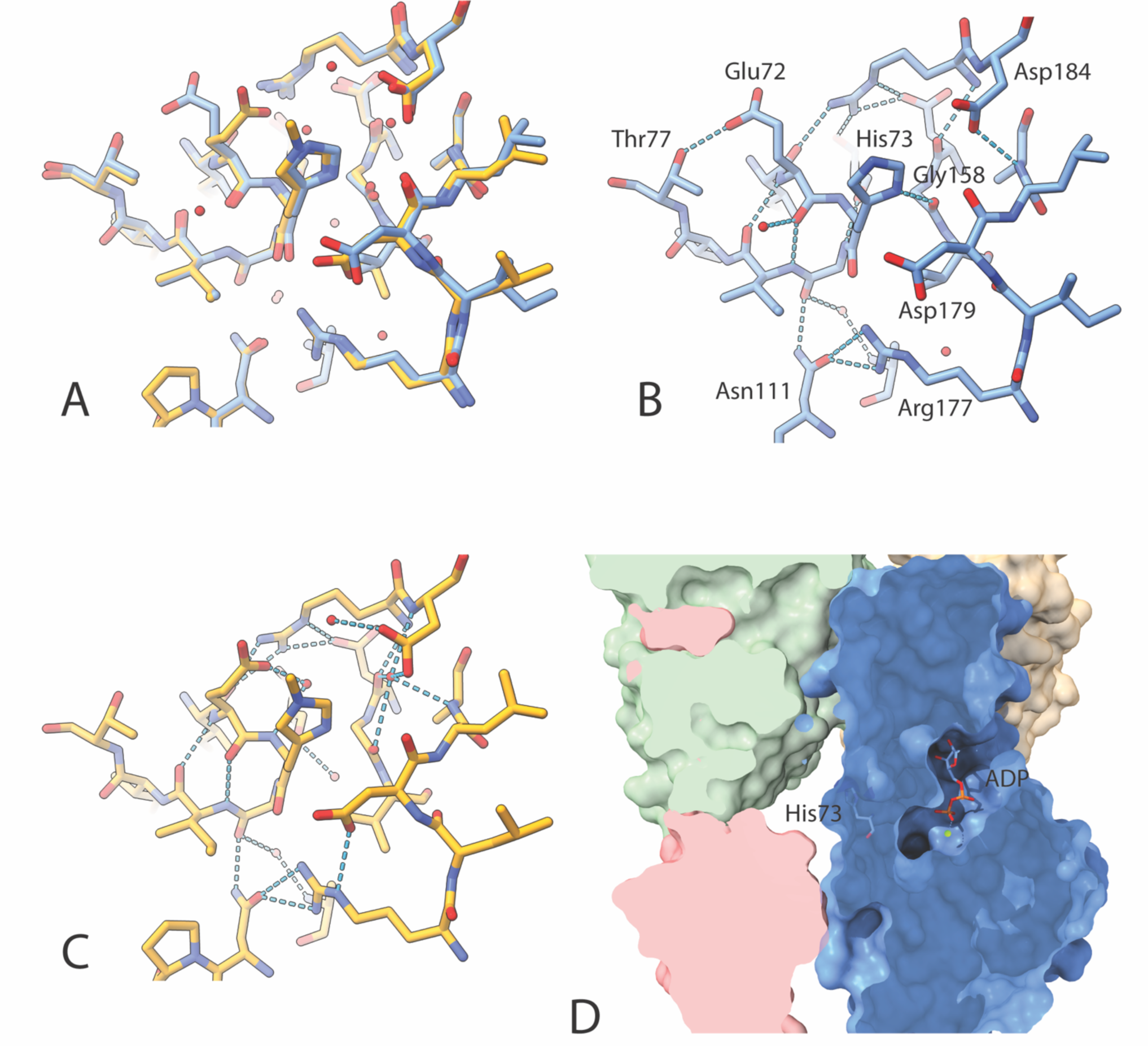
Comparison of yeast and mammalian actin in the vicinity of His73. (A) Alignment of the yeast (cornflower blue) and mammalian structures (orange; PDB ID:8A2T). (B) Yeast structure alone, with selected amino acids labeled and predicted H-bonds/salt bridges. (C) Equivalent view of the mammalian structure. (D) Cross section of the yeast actin filament showing the locations of His73 and ADP. (See also Fig. S3).

#### The W-loop, D-loop and H-loop

A crucial element of inter-subunit interactions in the longitudinal direction of the the filament is that between the W-loop (165-172) and D-loop (40-50) of respective subunits; the W-loop is a region of sequence divergence, with substitutions at positions 167,169 and 170 (Fig. 4). A water forms a central hub in an extended network in rabbit actin (Marked H_2_O in Fig. 4B) which appears to be absent in yeast actin as a result of substitutions at positions 167 and 292 (Fig. 4A,B). Thus, in yeast actin, the W loop is less constrained by the intra-subunit H-bond network (compare Fig. 4A with Fig. 4B) and can therefore shift its backbone towards the D-loop in the neighboring subunit (Note Ser170 in Fig. 4C), with the D-loop also shifting. The rearrangements in the vicinity of the W-loop in yeast actin enable a hydrogen bonding interaction with Gln49 of the D-loop in the neighboring subunit (Fig. 4). The overall impact of these changes is that the rabbit actin has two regions of extensive H-bonding in the vicinity of Asp292 and Tyr169 while the yeast actin has only one, in the region of Tyr169. This ‘loosening’ of the hydrogen bonding anchor of the W-loop could contribute to the apparent overall increased flexibility of yeast actin filaments^16,18^. Xu *et al.*, using a model-based sharpening method^39,17^, have attributed the stiffening of rabbit muscle F-actin to the presence of a coordinated Mg^2+^ ion indirectly bound to Glu167 via a water molecule. However, models of skeletal actin (e.g. PDB ID:8A2T^4^) which have sufficiently high resolution to reveal water molecules, do not indicate such an arrangement.

**Figure 4.**
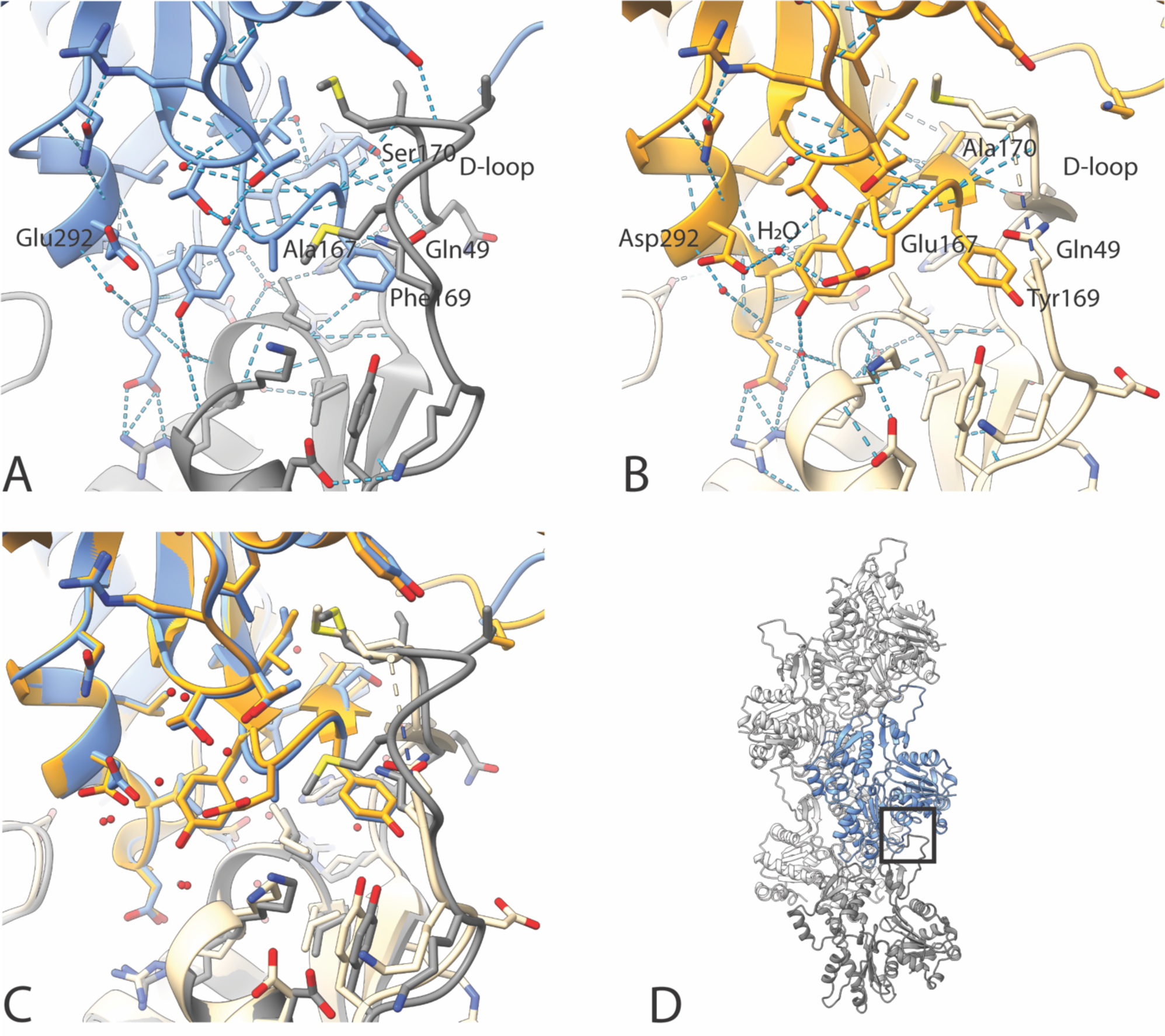
W-loop:D-loop interactions between subunits-comparison between yeast and mammalian F-actin. (A) Key interactions of the D-loop of one subunit (dark gray) and a neighboring subunit (including its W-loop) (blue) in yeast F-actin. (B) Equivalent interactions in mammalian actin (PDB ID:8A2T). (C) Structural alignments of yeast actin (blue/dark gray) and mammalian actin (orange/light gray). Waters are shown as small and large red spheres respectively. (See also Figs. S4-S5)

The difference in longitudinal contact strength was previously suggested to be due to the Glu167Ala substitution in yeast actin^40^. While it is the case that Ala167 in yeast does not participate in a salt bridge it is notable that Ser170 (Ala170 in skeletal muscle actin) facilitates additional H-bond interactions with the D-loop (Figure 4A,B). If this bonding network is disrupted in a Ser170Cys substitution in yeast, the polymerization rate is reduced^41^. These additional interactions may also account for the apparent increased ordering of the D-loop in yeast actin (Fig. S1B).

The D-loop is susceptible to proteolytic cleavage by the protease subtilisin between Met47 and Gly48 of the D-loop (Fig. S1B)^23^. The rate of subtilisin digestion of yeast F-actin was reported to be approximately 10-fold faster than muscle actin, and previously this was suggested to indicate greater flexibility compared with the D-loop of the skeletal muscle isoform^15^. The inter-residue bonding in the yeast F-actin structure however does not support the idea of greater flexibility in the D-loop region. Rather we would consider that the higher level of proteolysis observed under the reported conditions is possibly due to increased turnover rate of the filaments themselves with subtilisin cleaving the actin monomers following disassembly.

In addition to the W-loop, the D-loop also interacts with the C-terminus of the adjacent subunit (Fig. S4A). A notable feature is the Val43Ile substitution in yeast actin compared to skeletal actin. If Phe375 in the adjacent subunit were to maintain the position adopted in the skeletal actin, this would result in a steric clash; thus Phe375 adopts the more exposed position seen in yeast. This is consistent with fluorescent and phosphorescent labeling studies of Cys374 that indicated that the C-terminus of yeast F-actin is more flexible and more exposed to the surrounding environment than the C-terminus of skeletal muscle F-actin^22^. To avoid steric clash between Ile43 and Val139 in the adjacent subunit, the mainchain of the D-loop shifts relative to that in chicken actin (Fig. S4A). Notable also in this region are Arg39 and His40 which interact with both adjacent inter-and intra-strand protomers through a hydrogen bonded network of waters (Fig. S4B). Thus the D-loop serves a critical role in bringing together three separate protomers of the filament.

The H-loop is also a region of relatively high sequence divergence, with several substitutions between residues 262-274. The loop in yeast actin shows only minor shifts in the backbone relative to skeletal actin (Fig. S5).

In conclusion, using our 2.5 Å map of yeast actin, we have investigated whether the formerly reported differences between vertebrate skeletal muscle F-actin and yeast F-actin were consistent with our higher resolution structural data. In yeast actin, the more open nucleotide binding pocket and differences in the hydrogen bonding network in the vicinity of histidine 73 are likely to affect ATP hydrolysis and phosphate release; rearrangements of intersubunit binding interactions in the vicinity of the D-loop and W-loop are likely to contribute to the increased flexibility of yeast actin filaments. Given the accessible genetics of yeast, our high resolution structure of F-actin provides a platform for future functional interrogation.

## ACKNOWLEDGMENTS

We thank Dr. Julien Bergeron and Dr. Nora Cronin for help with data collection at the Francis Crick Institute; Professor Mohan Balasubramanian, Dr Tomo Hatano, Dr Andrejus Suchenko (University of Warwick) for support in expressing and purifying yeast actin, and Professor Bruce Goode, (Brandeis University) for valuable discussion. Local data collection and image processing was performed in the University of Sheffield, School of Biosciences EM facility, partly funded through the University of Sheffield Imagine: Imaging Life programme. KRA was supported by a Leverhulme Research Fellowship (RF-2021-099\2); SRS was supported by a Ph.D. studentship through the BBSRC White Rose Doctoral Training Programme (grant number BB/M011151/1). We thank Prof. Sarah Harris (University of Sheffield) for stimulating discussions on nucleotide binding.

## AUTHOR CONTRIBUTIONS

Conceptualization, K.R.A and P.A.B.; Methodology, K.R.A., P.A.B., S.J.S, S.B.T.and I.L.; Experimentation, S.J.S., S.B.T.and I.L.; Writing, K.R.A., P.A.B.; Visualization, P.A.B and K.R.A.; Supervision, K.R.A. and P.A.B.; Funding Acquisition, K.R.A. and P.A.B.

## SUPPLEMENTAL INFORMATION

Document S1. Figures S1–S5 and Table S1

## METHODS

### Data availability

EM maps and molecular structures generated in this study have been deposited at PDB and EMDB data banks under entry codes PDB ID 9GO5, EMD-51491, respectively.

### Yeast actin purification

The *Pichia pastoris* yeast strain used in this study to express *S.cerevisiae ACT1* was a kind gift from the Balasubramanian lab (University of Warwick). This strain is the X-33 *Pichia pastoris* strain (ThermoScientific) carrying an integrated plasmid (pPICZc-*Sc*Act1-Thy-β4-8His). This allowed expression of a thymosin beta-4-actin fusion. When this fusion protein is expressed, the Thyβ4 sequesters the recombinant G-actin, preventing interactions with both the barbed and pointed end of the monomer (Fig. 1C). Crucially, the bound thyβ4 makes the recombinant actin unavailable for polymerisation with the native pool of *Pichia* G-actin. Expression is under control of the AOX1 promoter allowing expression to be induced on addition of methanol to the growth medium. Cells were grown and induced to express *S. cerevisiae* actin as described^30^. Cells were broken using the freezer mill at the University of Warwick.

The actin was purified according to the published protocol with chymotrypsin cleavage being used to release the purified actin from the fusion as a chymotrypsin site is available immediately after the final actin residue Phe375^30^. Concentration of purified actin was estimated from actin standards on a Coomassie stained gel and for the experiments here the concentration of the preparation was 25 µM (approx. 10.5 mg/ml). Actin was kept in G-buffer: 10 mM Tris (pH 7.5), 0.2 mM CaCl2, 0.2 mM ATP (pH 7†), 0.5 mM DTT. Polymerisation competence was verified in a sedimentation assay following addition of polymerisation salts (KME) to a 50 µl assay volume containing 3 µM actin (KME - 50 mM KCl, 1 mM MgCl2, 1 mM EGTA, 10 mM Tris (pH 8.0). Samples were left for 1 hour at room temperature (20°C) prior to centrifugation at 90000 rpm in a Beckman ultracentrifuge for 15 minutes.

### Polymerisation of actin for CryoEM

Freshly prepared G-buffer was used to dilute G-actin giving a final sample volume of 20-30 μl. 10x KME polymerisation salts were added to 1x final. Polymerisation was allowed to occur at room temperature for approximately 1 hr.

Cryo grids were prepared using a Leica EM GP automatic plunge freezer (Leica Microsystems). An optimal concentration of 1 μM G-actin was used. At this concentration the filament density on the grid was high with limited filament overlap. 3 µl of actin was added to copper Quantifoil R2/2 300 mesh holey carbon grids. Excess liquid was blotted for 5 seconds from the mesh side of the grid (back blotting) and the grids were plunge-frozen in liquid ethane. Frozen grids were stored in liquid nitrogen until imaged.

### CryoEM data collection and image processing

CryoEM images were collected on a Titan Krios microscope (ThermoFisher Scientific) operated at 300kV and recorded on a K3 direct electron detector (Gatan Inc.) operated in super-resolution mode. The images were collected at a nominal magnification of 105,000X such that the object level pixel size was 0.834 Å/pixel (super-resolution pixel size of 0.417 Å/pixel). The images were recorded as 2.3 second movies divided into 27 frames. The total dose was 27 electron/Å^2^ and the fluence was 1 electron/Å^2^/frame.

All image processing jobs and three-dimensional (3D) reconstructions were performed using CryoSPARC (versions 3 and 4)^31^ (Structura Biotechnology Inc.). The individual superresolution movie frames were binned by 2 and the frames were aligned using alignparts_lmbfgs^42^ as implemented within Cryosparc. The contrast transfer function (CTF) of the aligned micrographs was estimated using the patch CTF routine of Cryosparc. After removing unsuitable micrographs (0.5 CTF fit resolution worse than 6 Å and average intensity higher than 3.8), 2540 micrographs were retained for further processing.

Particles were identified using template free filament tracer and extracted with a box size of 880 Å (the large box size allowed inclusion of two helical turns) and downscaled to a pixel size of 2 Å/pixel. At this stage 559,574 particles were selected. These particles were subjected to one round of 2D classification and 267,843 particles contributing to selected optimal 2D class averages (determined by visual inspection) were retained for further processing.

These particles were used to perform an initial helical reconstruction resulting in a 4 Å map. Using the particle orientation determined from the initial refinement as a starting point, the symmetry parameters were further refined. 2D templates were generated from this initial 3D map and used for a second round of template-based particle picking resulting in 2,058,503 particles. These particles were refined using the initial helical parameters as a starting point. resulting in a 2.8 Å map. This was followed by refinement of the defocus values of each particle and the beam tilt and higher order aberrations were estimated for each micrograph. Another round of helical refinement was performed using the updated CTF parameters. For this round the non-uniform refinement routine was used along with a finer search of the helical symmetry parameters providing the parameters listed in Table S1. This led to a map with a nominal resolution of 2.5 Å. In order to ensure that the reconstruction was not trapped in a local symmetry minimum, the refinement was repeated without applying helical symmetry and the resulting structure was identical to the one obtained with helical symmetry (data not shown).

The resolutions of the CryoEM maps were estimated from the gold standard Fourier Shell Correlation (FSC) curves^43^ calculated in CryoSPARC and are reported according to the 0.143 cutoff criterion. The FSC curves were corrected for the convolution effect of a soft mask applied to the half-maps using phase randomization^44^. To prevent overfitting during refinement, it was ensured that particles picked from the same filament were placed in the same half-set for gold standard FSC resolution estimation. The local resolution of the map was calculated in Cryosparc using the 0.143 FSC criterion.

### Model building and refinement

Coordinates for one subunit of rabbit F-actin (PDB ID:8A2T)^4^ were fitted into the sharpened CryoEM using the ‘Fit in Map’ tool in ChimeraX^45^. The fitted model was examined in Coot^46^, residues mutated to those of yeast where required and missing residues in the D-loop built in. Fits of side chain rotamers and main chain were optimized either by real space refinement within Coot or in ChimeraX/Isolde^47^.

Five copies of the subunit were fitted into the helical density of the CryoEM map and subjected to real space refinement within Phenix^48^, with non-crystallographic symmetry (NCS) constraints applied and refined. A central subunit of the refined structure was rebuilt where necessary to optimize geometry and clashes. A new filament model was built from this subunit using the refined symmetry operators, and subjected to further cycles of real space refinement and rebuilding.

Putative water molecules were added automatically in Coot with a 3 sigma density threshold around one central subunit. These were inspected manually and any found in noisy or highly asymmetric density were removed. NCS was applied to generate a five-subunit model and any clashing waters or waters with no clear hydrogen bonding interactions were removed. A final cycle of NCS constrained refinement in Phenix was performed with a final five-subunit filament model generated from the central refined protomer.

