## supplemental information for "CryoEM Reconstruction of Yeast ADP-Actin Filament at 2.5 Å resolution. A comparison with mammalian and avian F-actin"

**Table S1**

| | |
|---|---|
| **Data collection and processing** | |
| Microscope | Titan Krios |
| Voltage (kV) | 300 |
| Detector | Gatan K3 |
| Magnification | 105,000 |
| Exposure (electrons/Å$^2$) | 27 |
| Exposure rate (electrons/pixel/s) | 17 |
| Pixel size (Å) | 0.834 |
| Defocus range (µm) | -0.5 to -2.4 |
| Imposed symmetry | C1, 2.76 Å rise, -167.2° twist |
| Initial particles | 2,581,668 |
| Final particles | 1,368,572 |
| Map resolution (Å) | 2.5 |
| FSC threshold | 0.143 |
| Map sharpening B factor (Å$^2$) | -107.1 |
| **Model refinement** | |
| Model resolution (Å) | 2.5 |
| FSC threshold | 0.5 |
| Model composition | 5 actin monomers |
| Non-hydrogen atoms | 14,955 |
| Amino acids out of 375 | 6-375 |
| Ligands | 5 Mg2+, 5 ADP |
| Waters | 350 |
| B-factors (mean) | |
| - protein | 26 |
| - ligand | 13 |
| - water | 26 |
| RMS deviations | |
| - bond lengths (Å) | 0.005 |
| - bond angles (°) | 1.085 |
| Validation | |
| - molprobity score | 1.3 |
| - clashscore | 3.2 |
| - rotamer outliers (%) | 1.96 |
| Ramachandran plot | |
| - favoured (%) | 98.4 |
| - allowed (%) | 1.6 |
| - disallowed (%) | 0 |
| EMRinger score | 4.1 |

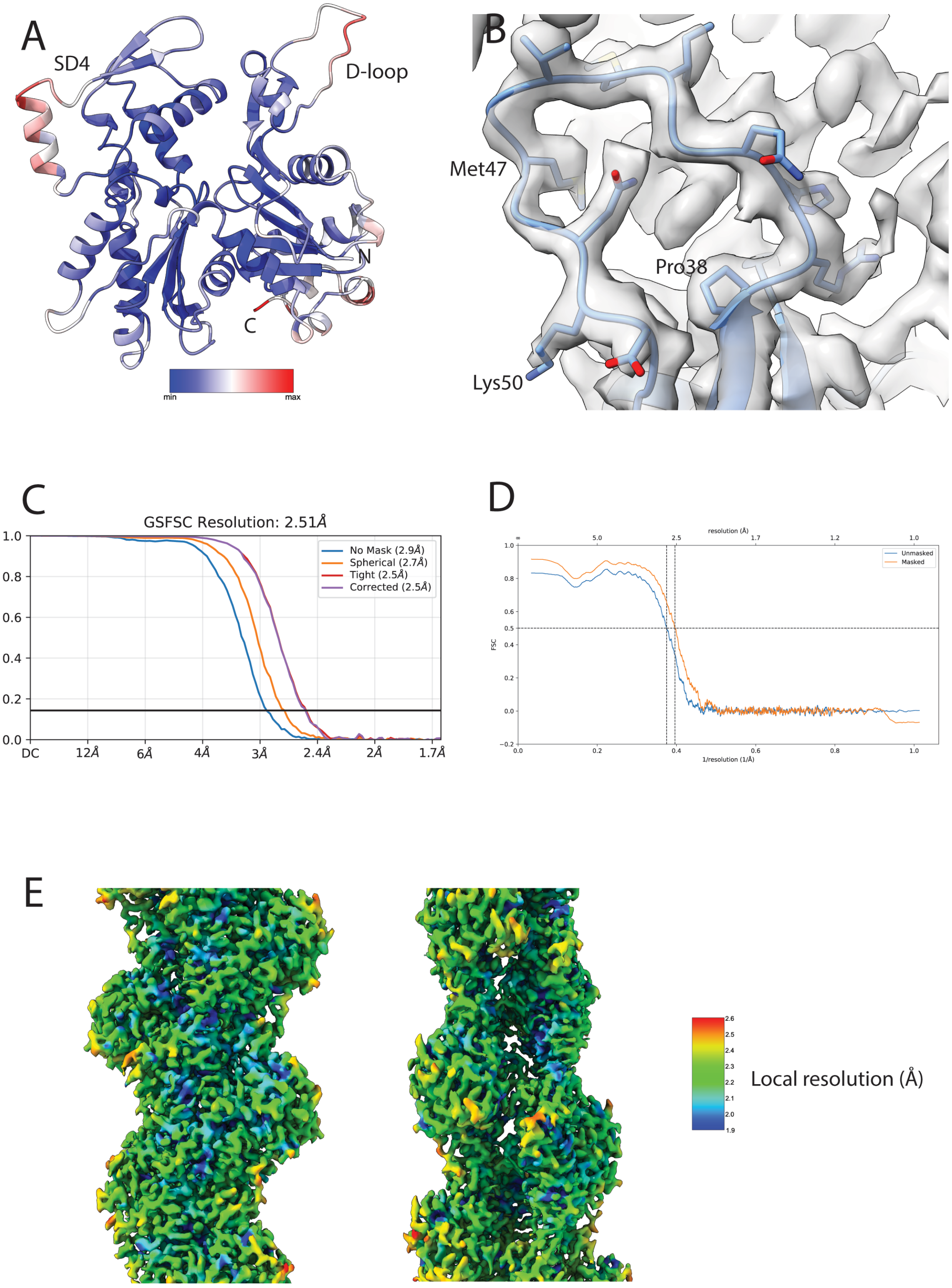

**Figure S1. Analysis of yeast F-actin CryoEM reconstruction, related to Fig. 1.**
(A) Yeast F-actin subunit fold colored by B-factor. (B) Clear density is resolved for the entire D-loop. (C) Plot of gold standard Fourier shell correlation against resolution for the yeast F-actin reconstruction. (D) Plot of Fourier shell correlation (FSC) between the yeast F-actin density map and the atomic model as a function of resolution. (E) Representation of local resolution estimates for the central promoters of the filament reconstruction. Two approximately orthogonal views are shown.

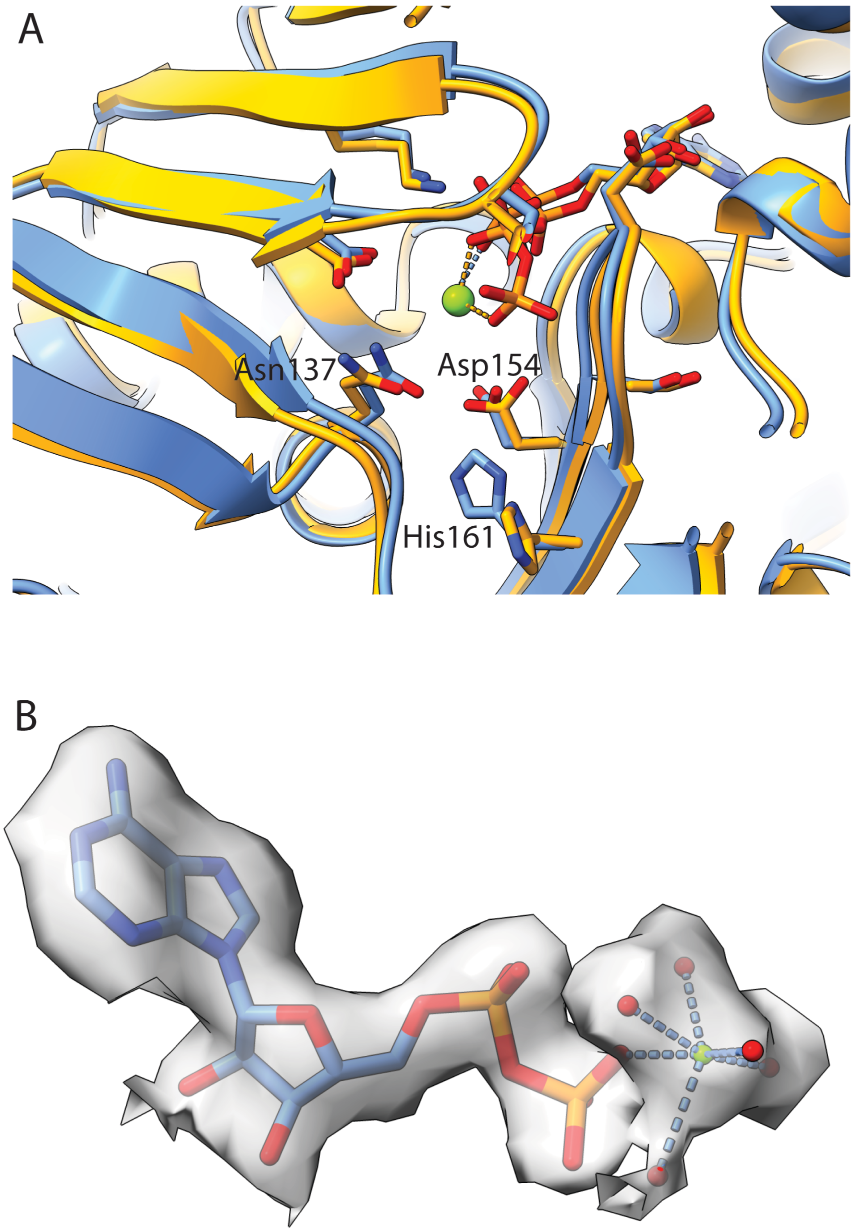

**Figure S2. The nucleotide binding site, related to Fig. 1, 2.**
(A) Alignment of residues in the yeast actin nucleotide binding site for ATP-bound G actin (orange) (PDB ID:1YAG, Vorobiev *et al.,* 2003) and ADP-bound F actin (blue). (B) Density of the CryoEM reconstruction around the ADP. Carbon- cornflour blue; nitrogen- dark blue; oxygen and water- red; phosphorous- orange; $Mg^{2+}$- green.

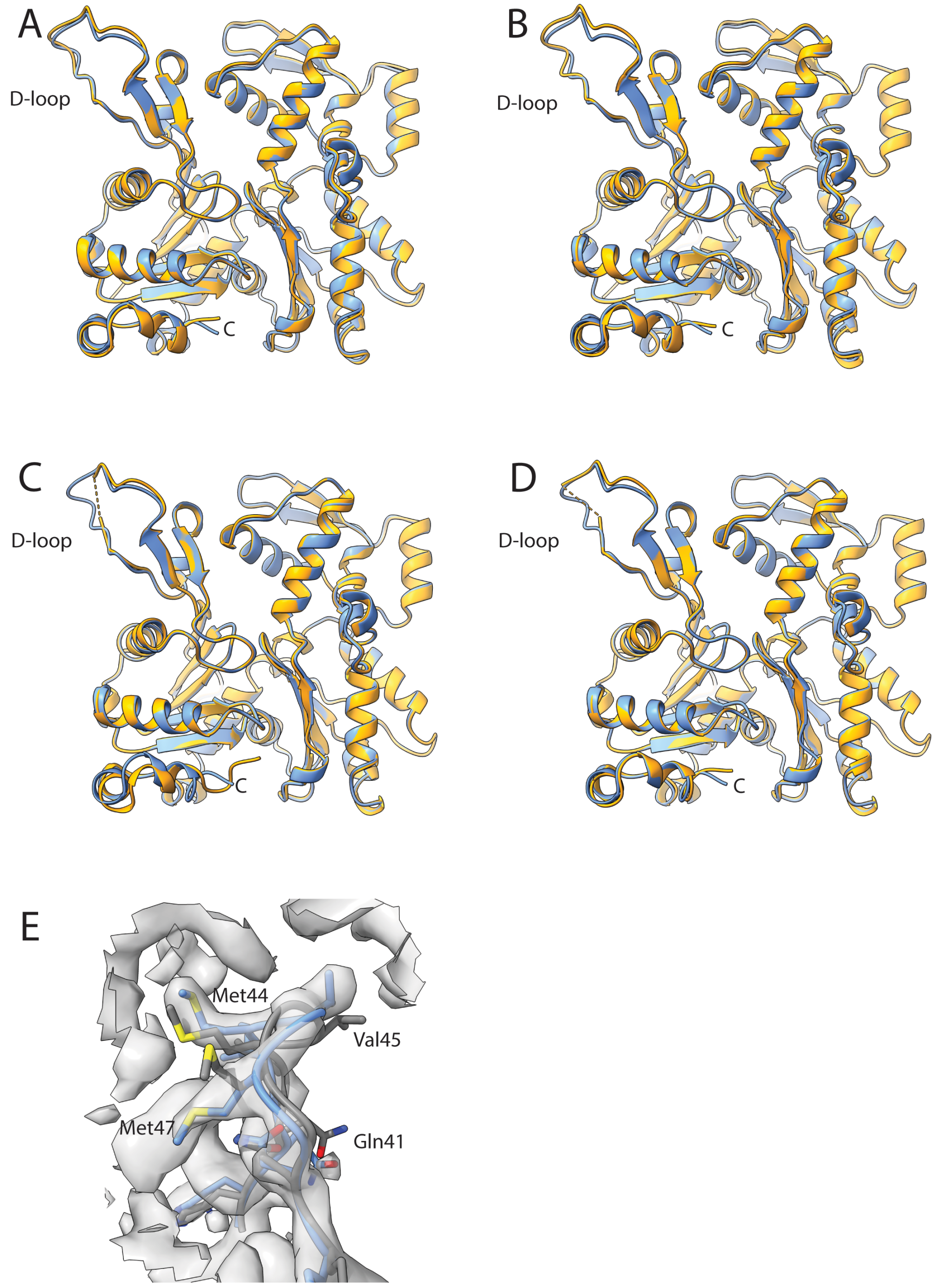

**Figure S3. Comparison of polypeptide backbone folds between yeast actin (blue) and vertebrate actin (orange), related to Figs. 3,4.**
Comparisons with (A) avian ADP-F-actin (PDB ID:8D13) (Reynolds *et al.,* 2022). (B) avian ADP-F-actin (PDB ID:7R8V) (Gong *et al.,* 2022). (C) mammalian ADP-F-actin (PDB ID:8A2T) (Oosterheert *et al.* 2022). (D) mammalian ADP-Pi-F-actin (8A2S) (Oosterheert *et al.* 2022). (E) Comparison of structure and CryoEM density in the vicinity of the D-loop of yeast ADP-F-actin at 2.5 Å (blue; current study) and that reported at 4.5 Å (gray) (PBD ID:8TI3) (Xu *et al*., 2024).

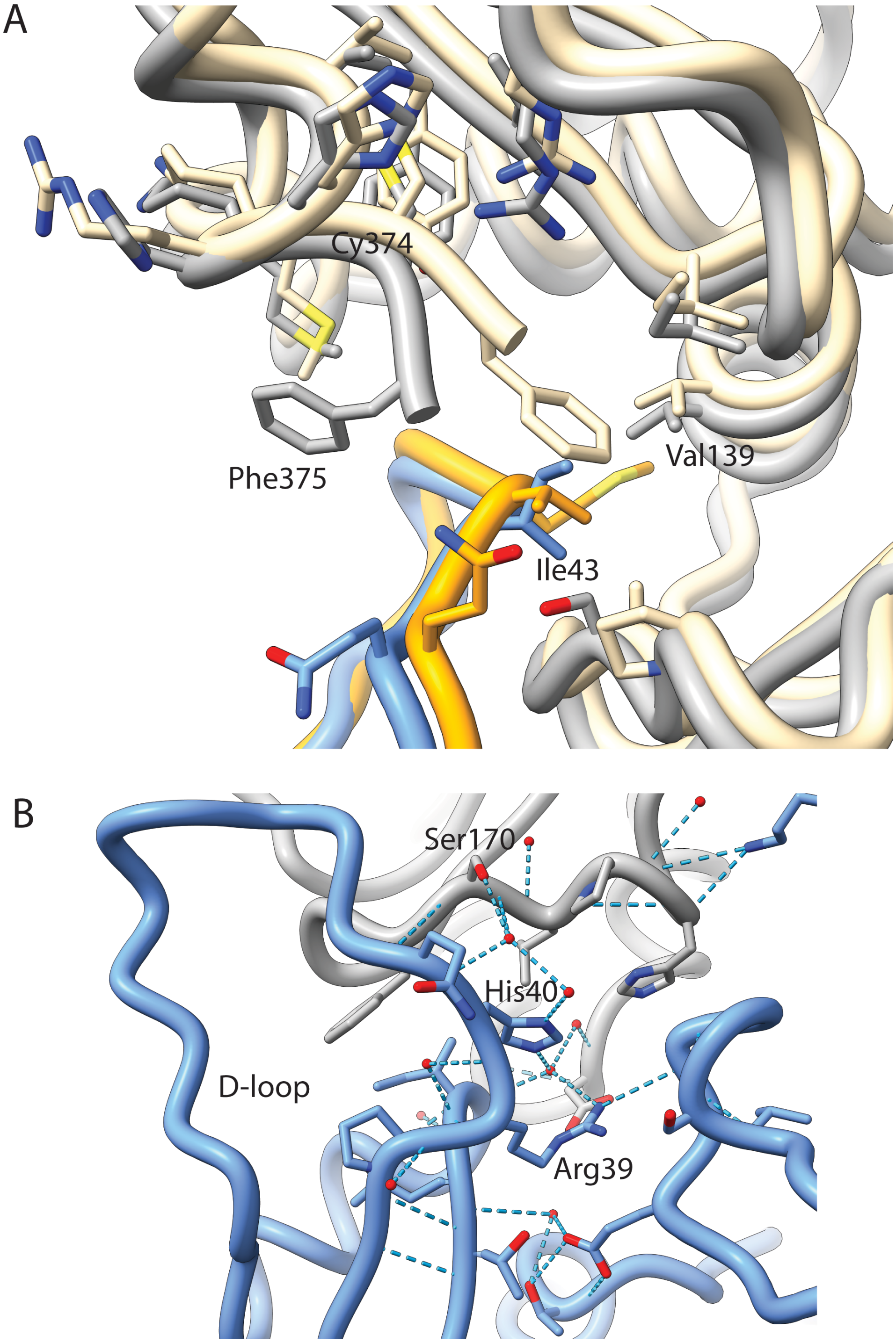

**Figure S4. Interactions with the D-loop in yeast and avian actin (PDB ID:8D13, Reynolds *et al.,* 2022), related to Fig. 4.**
(A) Comparison of interactions with the C-terminus of the neighboring subunit. Yeast subunits- blue and gray; avian subunits- orange and straw. (B) Predicted H-bonding network in the vicinity of His40 of the D-loop (blue) with the neighboring subunit (gray).

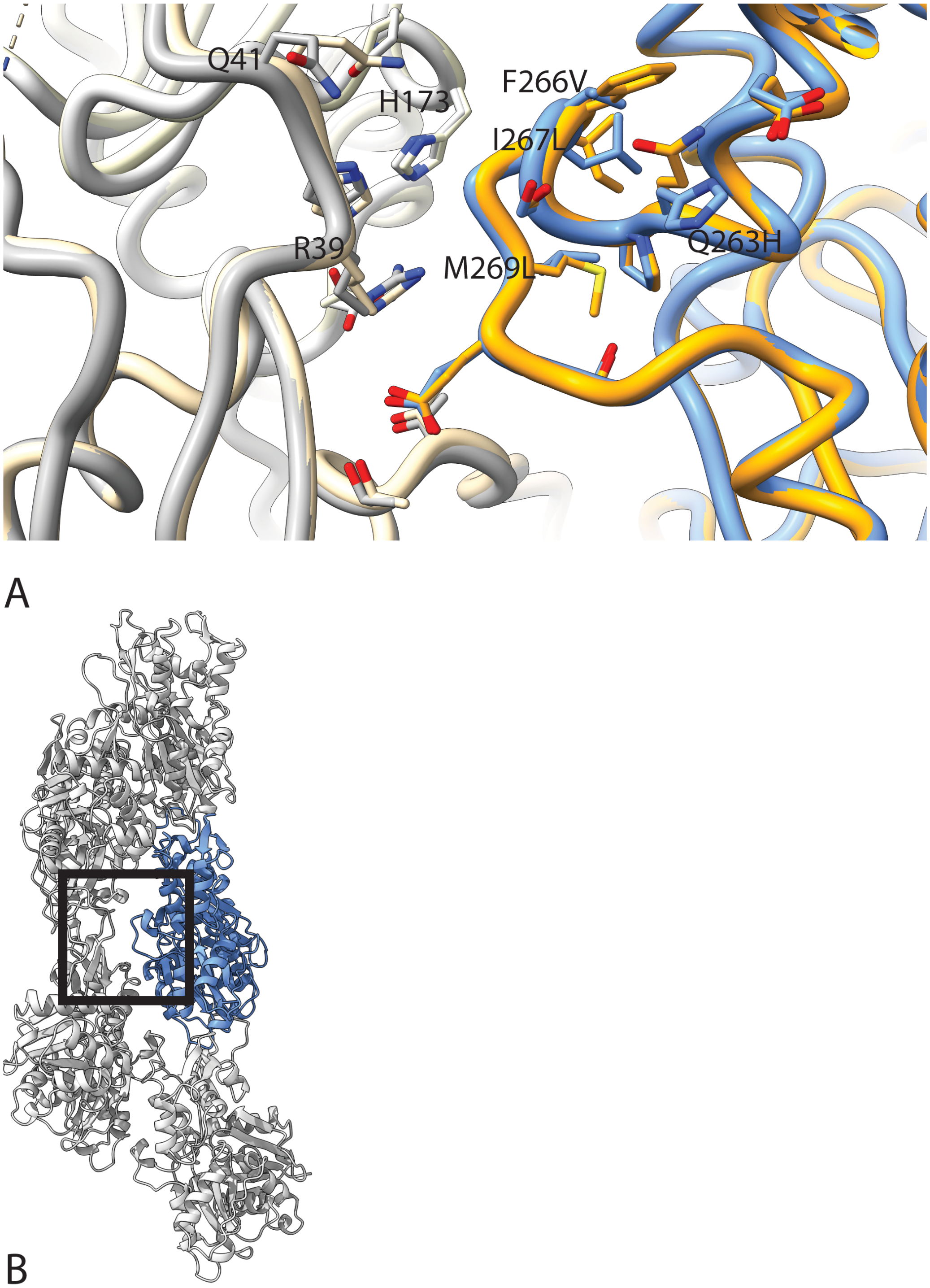

**Figure S5. Comparison of H-loop interactions in yeast and mammalian F-actin (PDB ID:8A2T, Oosterheert *et al.* 2022), related to Fig. 1**

(A) Selected residue substitutions (yeast versus mammalian actin) labeled for the H-loop for one yeast subunit (blue) and nearby residues in the adjacent strand (gray). For comparison, mammalian equivalents are shown in orange and straw. (B) Five central protomors of the yeast F actin, with one colored blue. The square box outlines the approximate region shown in (A).
